# Decontamination of ambient RNA in single-cell RNA-seq with DecontX

**DOI:** 10.1101/704015

**Authors:** Shiyi Yang, Sean E. Corbett, Yusuke Koga, Zhe Wang, W. Evan Johnson, Masanao Yajima, Joshua D. Campbell

## Abstract

Droplet-based microfluidic devices have become widely used to perform single-cell RNA sequencing (scRNA-seq) and discover novel cellular heterogeneity in complex biological systems. However, ambient RNA present in the cell suspension can be incorporated into these droplets and aberrantly counted along with a cell’s native mRNA. This results in cross-contamination of transcripts between different cell populations and can potentially decrease the precision of downstream analyses. We developed a novel hierarchical Bayesian method called DecontX to estimate and remove contamination in individual cells from scRNA-seq data. DecontX accurately predicted the proportion of contaminated counts in a mixture of mouse and human cells. Decontamination of PBMC datasets removed aberrant expression of cell type specific marker genes from other cell types and improved overall separation of cell clusters. In general, DecontX can be incorporated into scRNA-seq workflows to assess quality of dissociation protocols and improve downstream analyses.

## INTRODUCTION

Single-cell RNA sequencing (scRNA-seq) has emerged as a powerful technique to study complex biological systems at single-cell resolution (Wang et al,2015). Droplet-based scRNA-seq platforms have been widely adopted because of their ability to profile a large number of cells at relatively low cost (Ziegenhain et al. 2017). These devices work by using droplets to partition cells into nanoliter reaction chambers along with beads harboring oligonucleotide primers with unique barcodes. Within each droplet, cells are lysed and the mRNAs will be tagged with the oligonucleortide primers to create barcoded cDNA after reverse transcription (Macosko et al,2015, Zilionis et al,2016, Zheng et al,2017).

Despite their many advantages, droplet-based single-cell technologies can suffer from the presence of cross-contamination from ambient RNA in each droplet. Ambient RNA is the pool of mRNA molecules that have been released in the cell suspension, likely from cells that are stressed or have undergone apoptosis. Cross-contamination occurs when the ambient RNA gets incorporated into the droplets and is barcoded and amplified along with a cell’s native mRNA (**Figure 1A**). Contamination from ambient RNA is evident when highly-expressed cell-type specific genes are observed at low levels in other cell populations. Different levels of contamination can be found in different droplets depending on the amount of ambient and native mRNA present. Two major goals of many scRNA-seq studies are to cluster cells into subpopulations and identify unique combinations of marker genes that define each cell population (Trapnell,2015). Ambient RNA can hinder these tasks by causing different cell populations to blend together and the expression of true marker genes to be detected across multiple cell populations. We developed a computational method called DecontX to estimate and remove ambient RNA for scRNA-seq data. We applied DecontX to three datasets to demonstrate its ability to accurately quantify and remove contamination within each cell from other populations and to improve downstream clustering.

**Figure 1:**
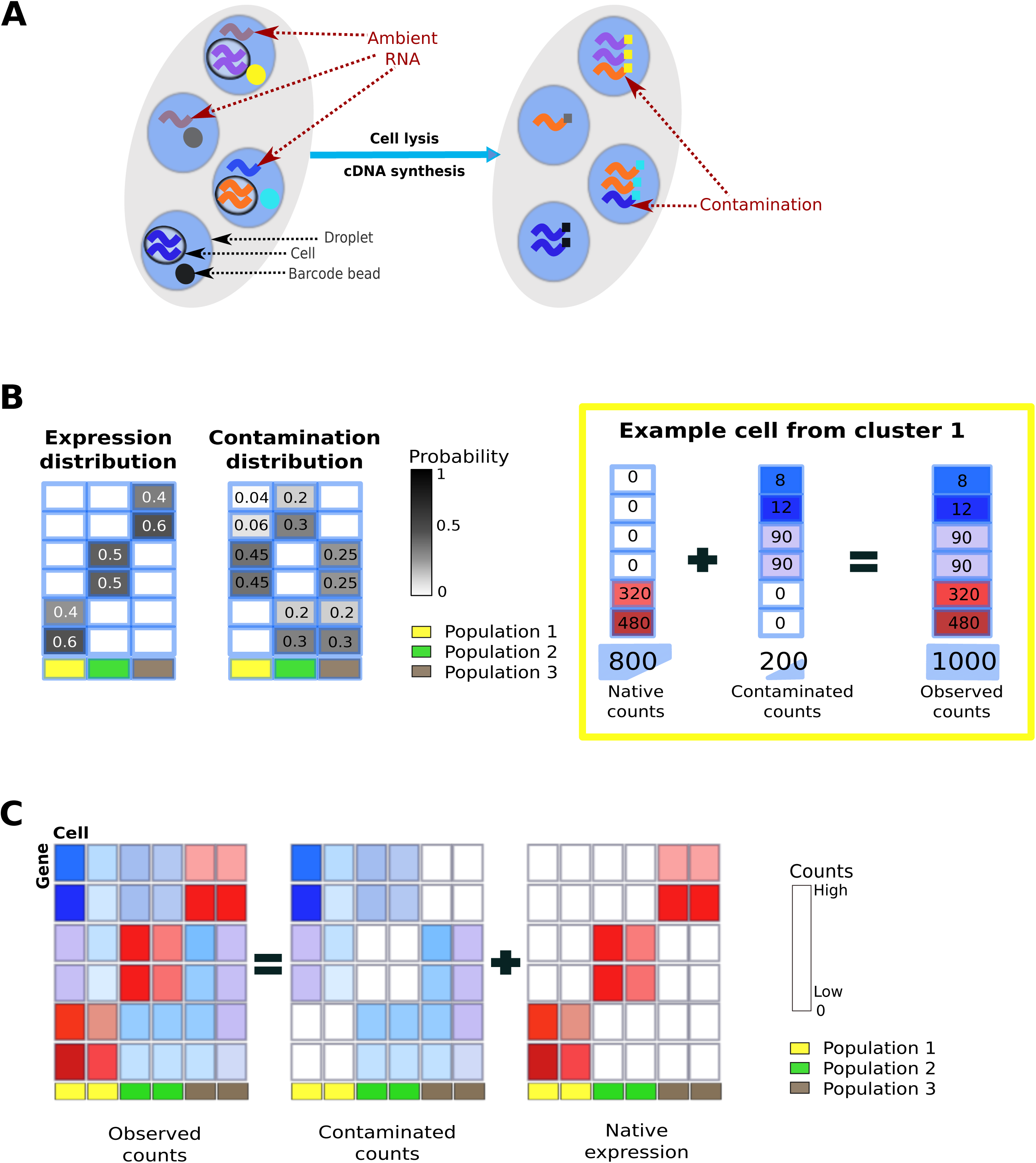
Overview of decontamination with DecontX. **(A)** In droplet-based microfluidic devices, ambient RNA can be incorporated into droplets along with oligonucleotide-barcoded beads and cells. Both native mRNA from the cell and contaminating ambient RNA will be barcoded and counted within a droplet. **(B) Left**: DecontX assumes that each cell is a mixture of two multinomial distributions: 1) a distribution of native transcripts from the cells true population and 2) a distribution of contaminating transcripts from all other cell populations captured in the assay. **Right**: Simulation of an example cell with 20% contamination. The 800 native transcripts are from the multinomial distribution for cell population 1 while the 200 contaminating transcripts are derived from a probability distribution that is a weighted combination of the two other populations. **(C)** DecontX will take an expression count matrix and cell cluster labels and estimate matrices of native expression and contamination from ambient RNA.

## RESULTS

To address the issue of contamination, we developed a novel Bayesian method called DecontX that identifies and removes contamination in individual cells. We assume the observed expression of a cell is a mixture of counts from two multinomial distributions: 1) a distribution of native transcript counts from the cell’s actual population and 2) a distribution of contaminating transcript counts from all other cell populations captured in the assay (**Online Methods**, **Figure 1B**). The native expression distribution for each cell population is characterized by a multinomial parameter *ϕ*_*k*_, where *ϕ*_*kg*_ is the probability of gene *g* being expressed in population *k*. Likewise, the contamination distribution for each cell population k is characterized by a multinomial parameter *η*_*k*_, where *η*_*kg*_ is the probability of gene *g* contaminating population *k*. Each individual cell *j* has a parameter *θ*_*j*_, which follows a beta distribution and represents the proportion of counts derived from the native expression distribution. Each transcript count has a hidden state, *y*_*jt*_, which follows a bernoulli distribution parameterized by *ϕ*_*j*_ and denotes the transcript’s membership to the native expression distribution (*y*_*jt*_ = 1) or contamination distribution (*y*_*jt*_ = 0). This framework is similar to a discrete Bayesian hierarchical model called latent Dirichlet allocation (LDA) (Blei et al,2003) where documents are mixtures of *K* topics and each topic is a mixture of words from a predefined vocabulary. However, rather than having *K* different distributions to model the mixtures of counts from different cell populations within each cell, we explicitly define the contamination distribution to be a weighted combination of all other cell population distributions. We use variational inference (Jordan et al,1999) to approximate posterior distributions to allow fast and scalable inference in large datasets (Blei et al,2017). Ultimately, DecontX will deconvolute a gene-by-cell count matrix and a vector of cell population labels into a matrix of contamination counts and a matrix of native counts which can be used in downstream analyses (**Figure 1C**).

To demonstrate the accuracy of DecontX, we utilized a public datase containing a mixture of fresh frozen human embryonic cells (HEK293T) and mouse embryonic fibroblasts (NIH3T3) cells from 10X Genomics. Using CellRanger (Zheng et al,2017), reads were uniquely aligned to a combined human-mouse reference genome (hg19 and mm10) to ensure that only reads specific to each organism will be counted while those that align to the genome of both organisms will be excluded. Cells were classified as human, mouse, or multiplets based on the levels of the organism-specific transcript counts (**Supplementary Figure 1**). The cells predicted to be either mouse or human still exhibited low levels of expression of counts aligning specifically to the other organism (**Figure 2A**). The proportion of mouse-specific genes in human cells was highly correlated to the distribution of expression in an average mouse cell (R=0.96; **Figure 2B**). Conversely, the proportion of human-specific genes in mouse cells was highly correlated to the distribution of expression in an average human cell (R=0.99; **Figure 2C**). These results also show that highly expressed genes in one cell subpopulation are more likely to contribute to contamination in other cell populations. Furthermore, while the median contamination was relatively low (1.09% in human cells and 2.75% in mouse cells), the percentage of contamination varied substantially from cell to cell (0.43%-45.09% in human; 1.25%-44.43% in mouse; **Figure 2D**) and demonstrates the need to have individual estimates of contamination for each cell.

**Figure 2:**
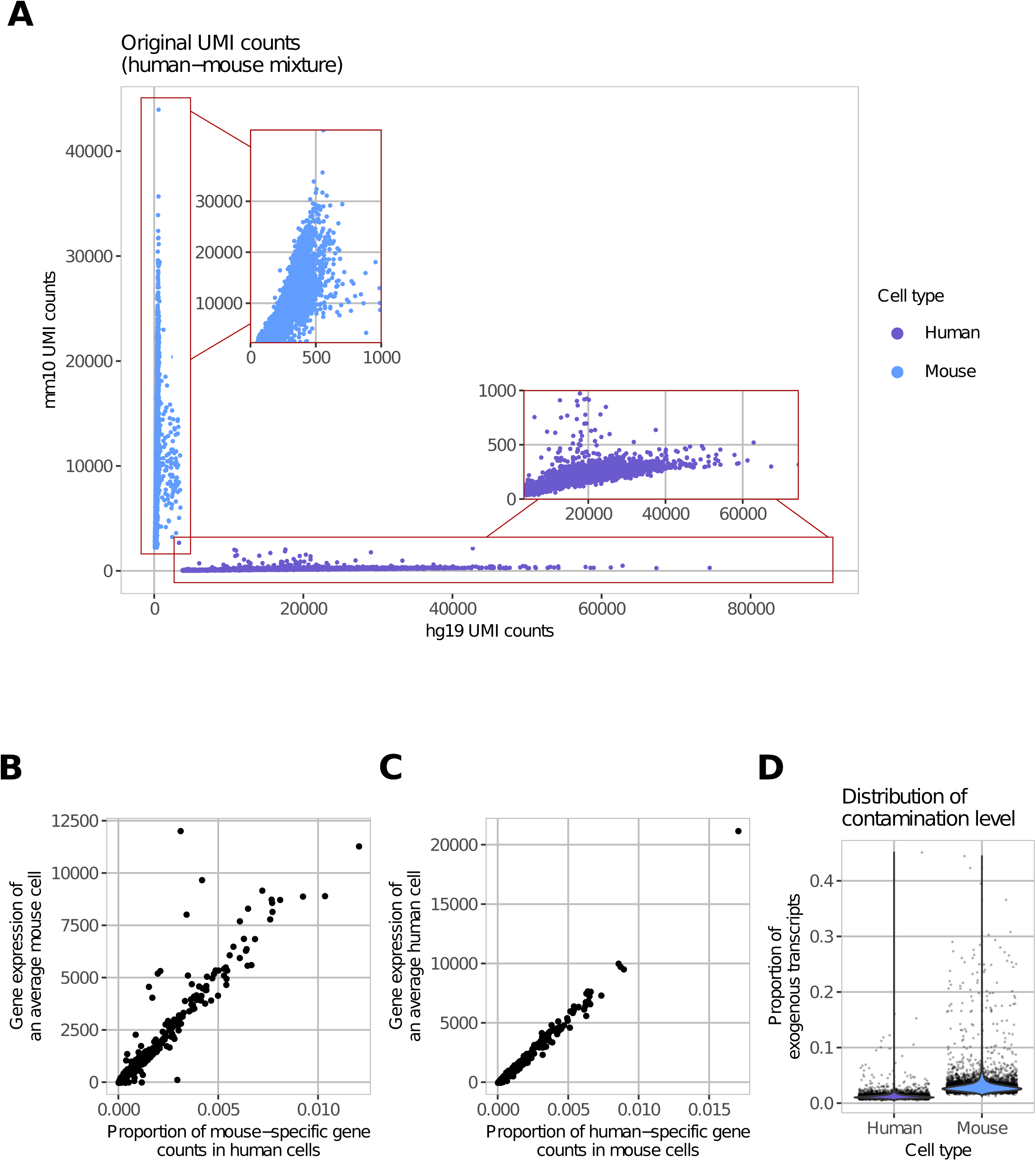
Contamination in a human-mouse cell mixture dataset. **(A)** The total number of UMIs aligned specifically to the mouse or human genome is plotted for each droplet. **(B)** The proportion of counts for mouse genes in human cells is highly correlated to the average expression of these genes across all mouse cells indicating that the amount of contamination for each gene is proportional to how highly that gene is expressed in the contaminating cell population. **(C)** Similarly, the proportion of counts for human genes in the mouse cells is highly correlated to the average expression of those genes across all human cells. **(D)** While each droplet is predicted to contain a single cell, the median percentage of contamination for human and mouse cells is 1.09% and 2.75%, respectively. The range of contamination is 0.43% - 45.09%indicating the need for contamination estimation for each individual cell.

We applied DecontX to 12,079 non-multiplet cells in the human-mouse mixture dataset. Most of the exogenous transcripts were identified and removed by DecontX (**Figure 3A**). The estimated proportion of contamination in individual human cells was highly correlated to the proportion of mouse-specific transcripts in those cells (R= 0.99; RMSE=0.002; **Figure 3B**). A high correlation was also observed in mouse cells (R=0.99; RMSE=0.005; **Figure 3C**), demonstrating the ability of DecontX to accurately detect contamination from other cell populations. The estimated gene-level contamination distributions for human or mouse cell populations were also highly correlated to the expression of an average mouse or human cell, respectively (**Supplementary Figure 2**).

**Figure 3:**
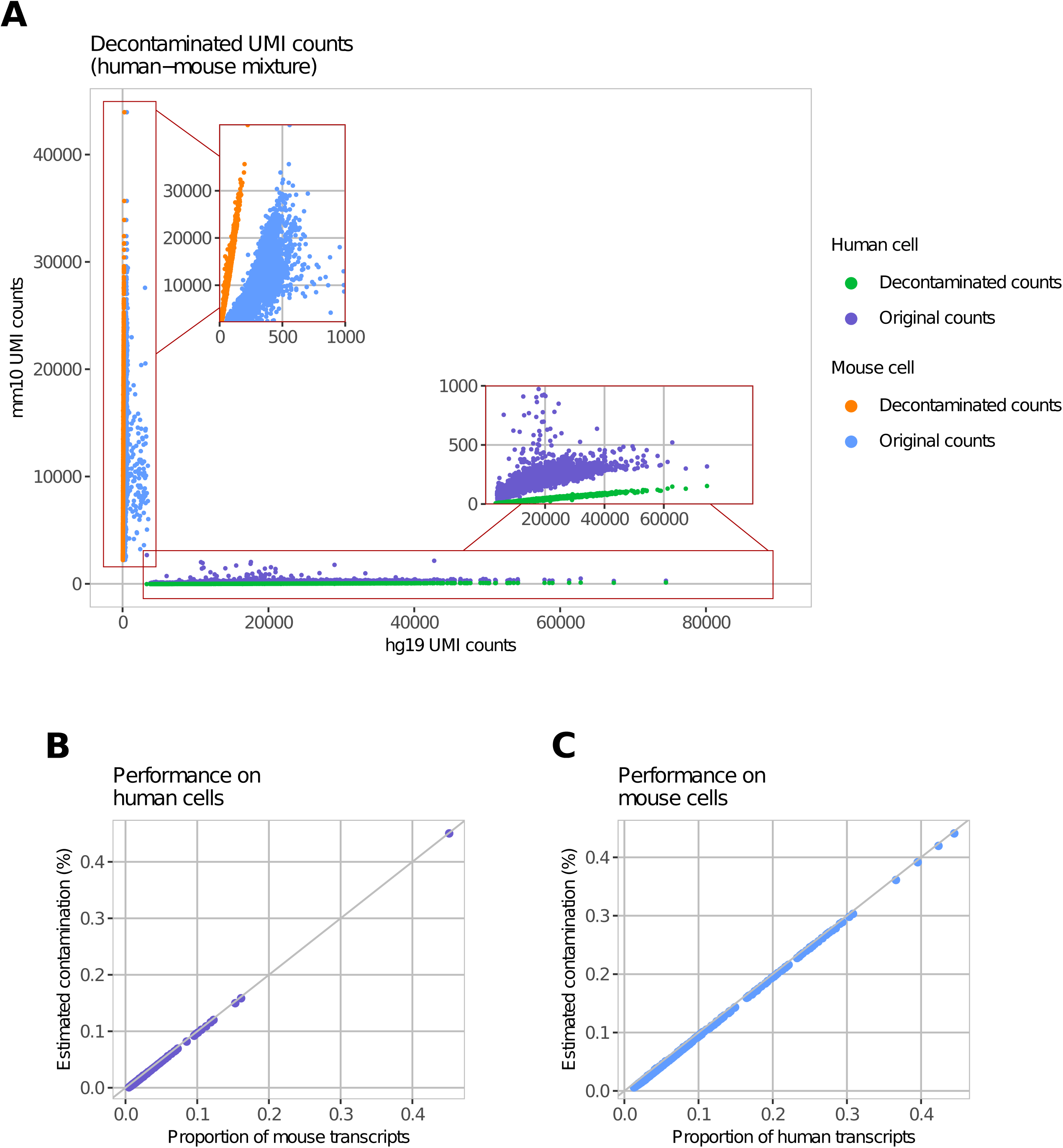
Decontamination of the human-mouse cell mixture dataset. **(A)** The number of human UMIs is again plotted against the number of mouse UMIs for each droplet before and after decontamination with DecontX. After DecontX, the median percentage of contaminating counts for each droplet is 0.26% (0.12% - 0.73%). **(B**, **C)** The DecontX-estimated contamination proportion is highly correlated to the known proportion of exogenous transcripts for each droplet predicted to have a human or mouse cell.

We next sought to understand the e?ect and extent of contamination in publicly-available scRNA-seq datasets of peripheral mononuclear cells (PBMCs). To establish baseline expression of cell-type specific marker genes in a settting with limited possibility for contamination, we examined 4 different immune populations (sorted PBMCs) isolated by flow cytometry and profiled with the 10X Genomics Chromiumin separate channels (Zheng et al, 2017). As each population was isolated and profiled in a different channel, gene markers for a specific immune population were detected at relatively low levels in other populations. For example, the mRNA expression of T-cell specific genes such as CD3E and CD3D were only found 0.07% in the B-cells sorted on CD20. Conversely, B-cell specific markers such as CD79A, CD79B and MS4A1 were only detected in 9.09% of T-cells sorted on CD8A or CD4 (**Figure 4A, 4B**). Similarly, low percentages of marker genes of other cell types could be found for B-cells and monocytes, monocytes and T-cells, T-cells and NK-cells, NK-cells and B-cells, and NK-cells and monoctyes (**Figure 4B**, **Supplementary Figure 3**).

**Figure 4:**
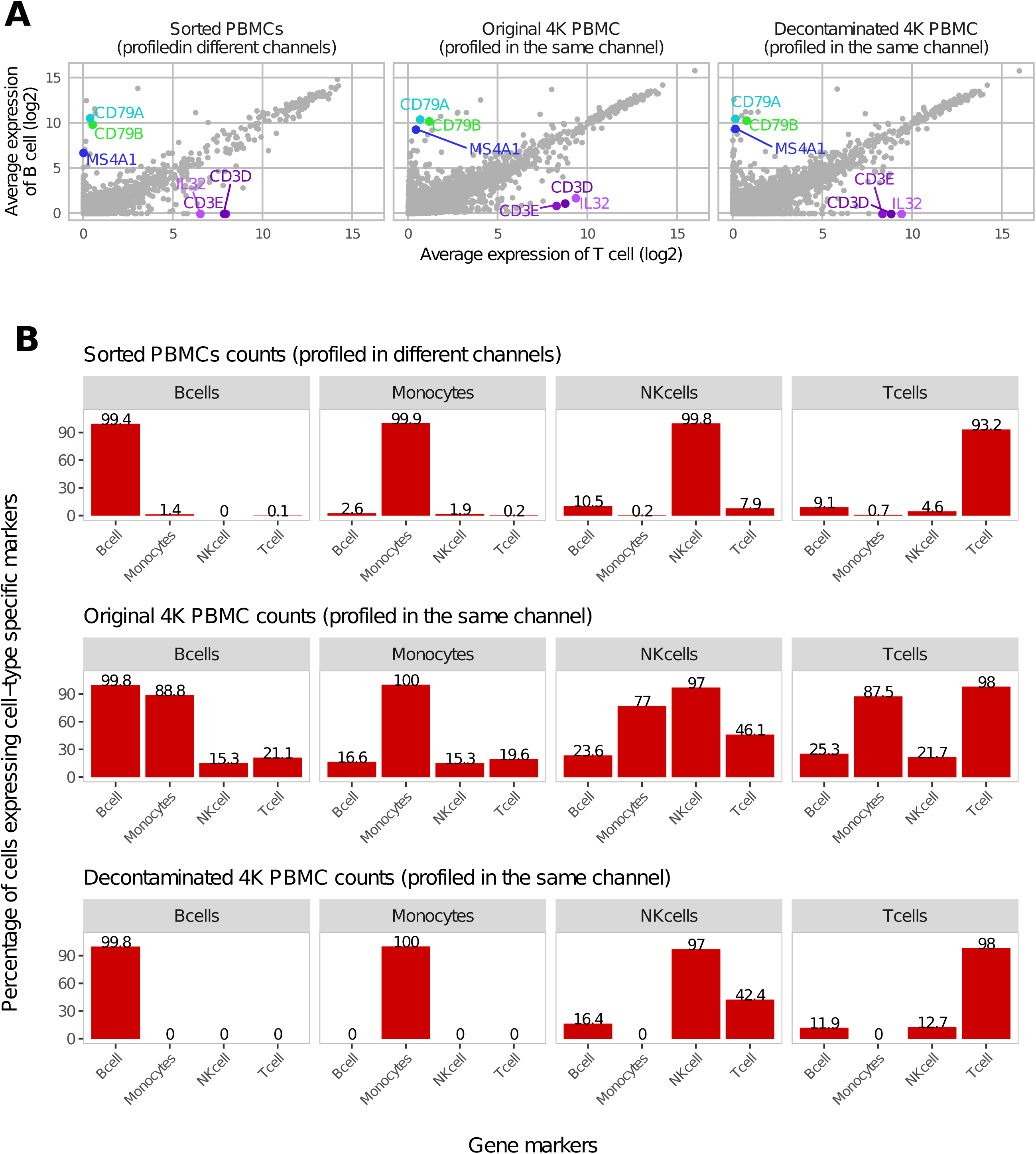
Expression of cell-type specific marker genes before and after decontamination in PBMCs. **(A)** For each gene, the average expression in the B-cell clusters is plotted against the average expression in T-cell clusters for three different datasets: data from sorted PMBCs profiled in different channels **(left)**; data from the PBMC 4K before decontamination **(middle)**; and the PBMC 4K data after decontamination with DecontX **(right). (B)** Percentage of cells expressing specific marker genes for different cell-types for three different datasets. Markers included CD79A, CD79B and MS4A1 for B-cells, CD3E and CD3D for T-cells, GNLY for NK-cells; and LYZ, S100A8 and S100A9 for monocytes.

In the second dataset, over four thousand PBMCs (4K PBMC) were isolated and profiled in a single channel of the 10X Genomics Chromium. Since cluster labels were not available from flow cytometry, we utilized Celda (Corbett et al,2019) to identify 19 cell populations where each population was a unique combination of 150 gene modules (**Supplementary Figures 5**, **6**). In contrast to the previous dataset, higher levels of cell-type specific marker genes could be detected in other cell types including CD3E and CD3D in 21.12% B-cell population; CD79A, CD79B and MS4A1 in 25.32% T-cell population (**Figure 4**); Likewise, higher level of a marker gene (GNLY) for NK-cells was found in monocytes and B-cells, marker genes (LYZ, S100A8 and S100A9) for monocytes in NK-cells, B-cells and T-cells (**Figure 4B**, **Supplementary Figure 3**). After we applied DecontX to remove contamination, the expression of T-cell specific marker genes was eliminated in B-cells and expression of B-cell specific marker genes was eliminated in T-cells (**Figure 4A**). The percentage of cells within each subpopulation that had expression of marker genes from other cell types markedly decreased (**Figure 4B**, **Supplementary Figure 3**). While the overall levels of the NK-cell marker GNLY was substantially reduced in T-cells and the T-cell markers CD3D and CD3E were reduced in NK-cells, some expression of these markers still remained in both populations. Decontaminated counts resulted in improved separation in two dimensions when applying tSNE (Maaten et al,2008) (**Figure 5A, 5B**). Additionally, the mean silhouette width, a measure of cluster stability and separation, improved from 0.04 on original normalized expression to 0.06 on normalized expression after decontamination (**Figure 5C**). Specifically, all clusters have improved mean silhouette width except for the cluster 17, which shows a decrease of average silhouette width from −0.002 to −0.042 (**Figure 5C**). Interestingly, cells from cluster 17 were predicted to be doublets by a doublet prediction method Scrublet (Wolock et al,2019) (**Figure 6A**). Cells predicted to be doublets by Scrublet are associated with higher contamination estimated by DecontX (p-value *<* 2e-16, **Figure 6**).In fact, all cells estimated to have high levels of contamination (*>* 70%) were predicted to be doublets by Scrublet suggesting that DecontX contamination estimates can be used as orthogonal information for doublet detection (**Figure 6C**).

**Figure 5:**
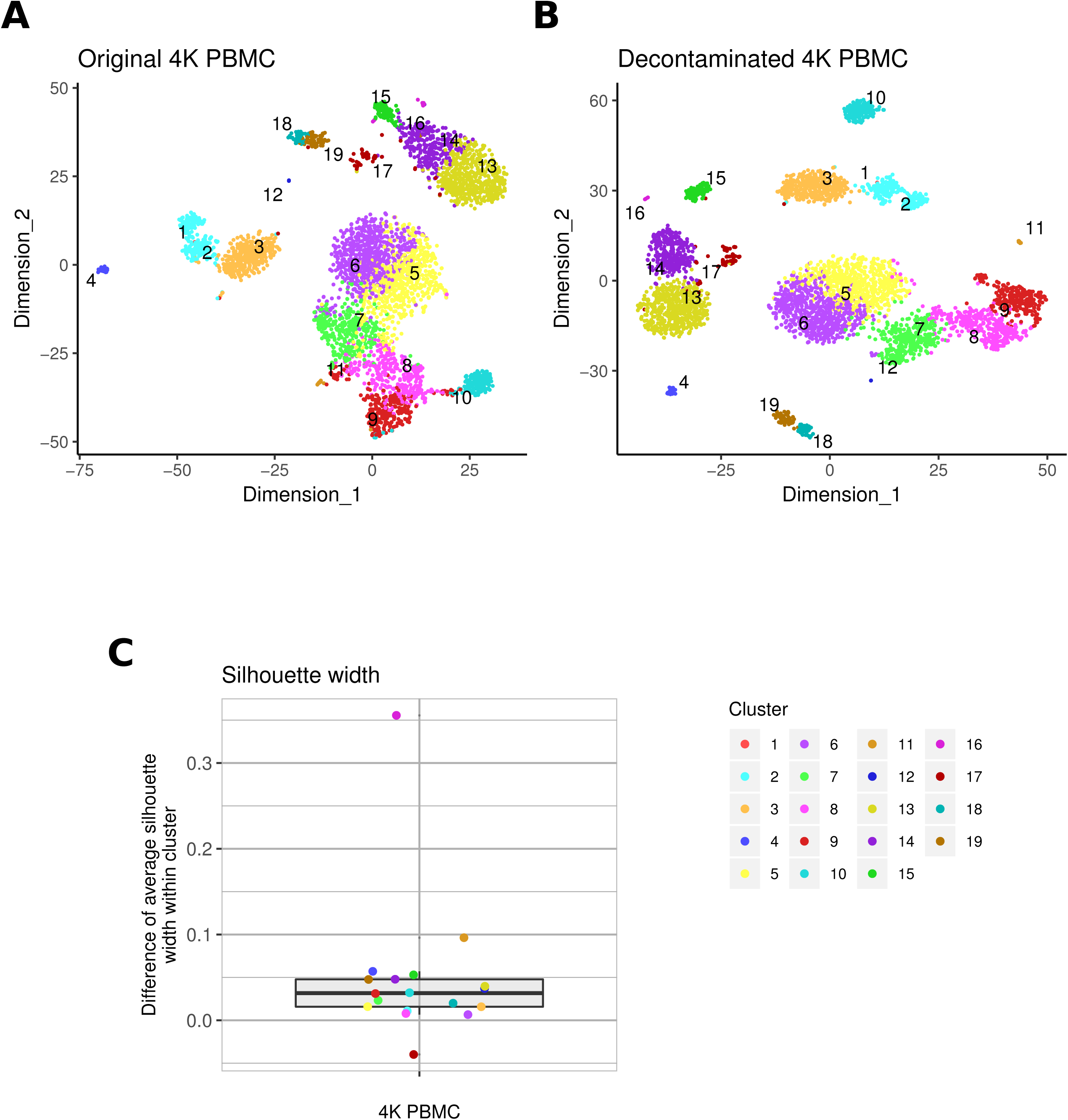
Cluster similarity before and after decontamination. **(A)** tSNE of 19 cell clusters from the PBMC 4K dataset before decontamination. **(B)** Decontamination with DecontX improved separation on tSNE between different cell clusters. **(C)** The mean silhouette width was derived for each cluster before and after decontamination with DecontX. Each point represents the difference in the mean silhouette width for each cluster. All clusters except 17 showed an increase in silhouette width after decontamination. Cluster 17 was predicted to contain mostly doublets by Scrublet. Cluster 1 had only one cell and was not included in the analysis.

**Figure 6:**
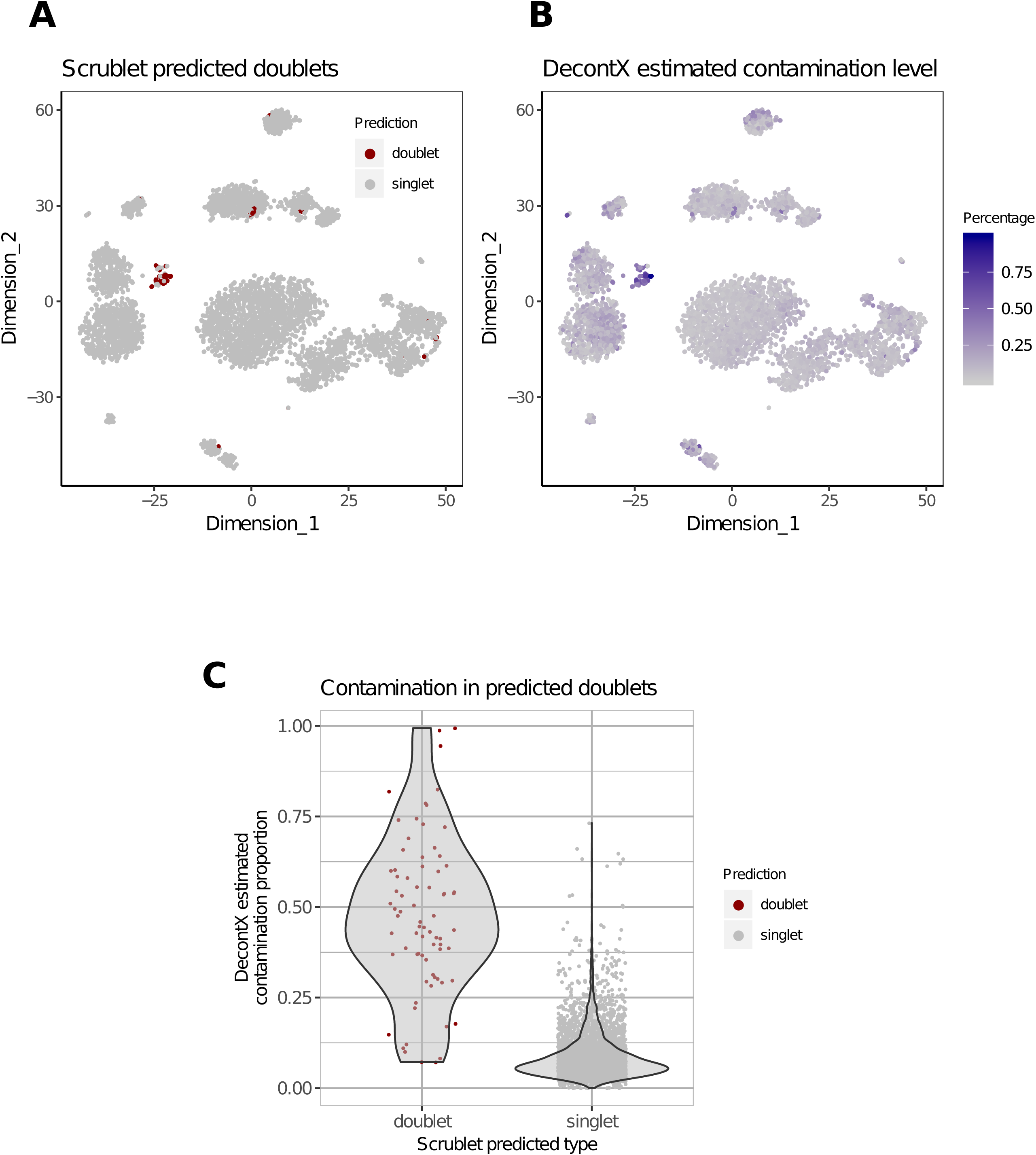
Comparison of contamination levels with predicted doublets. **(A)** tSNE of DecontX decontaminated PBMC 4K data. Red cells are predicted doublets by Scrublet. **(B)** Each cell is colored by contamination level estimated by DecontX. **(C)** Predicted doublets had significantly higher levels of estimated contamination compared to singlets. The median contamination for doublets was 45.95% (7.18% - 99.39%) while the median for singlets was 6.81% (0.02% - 73.18%).

## DISCUSSION

We developed a method called DecontX to estimate the percentage of cross contamination within each cell due to ambient RNA in droplet-based single-cell RNA sequencing experiments. In human-mouse mixture data, DecontX was able to accurately estimate the percentage of exogenous transcripts. After estimating and removing contaminated transcripts in 4K PBMC data, the profiles of key marker genes for each subpopulation better resembled those from sorted PBMCs. Furthermore, decontamination resulted in improved downstream clustering and visualization. The cells estimated to be highly contaminated by DecontX in 4K PBMC were also estimated to be doublets by Scrublet. Therefore, high contamination levels may also be useful as a quality control criterion for excluding cells. Additionally, estimating the levels of background RNA contributing to the contamination will be important for quality assessment of cell dissociation protocols.

By utilizing raw counts for estimation of the multinomial distributions, DecontX eliminates the potential variability that could be introduced by different normalization methods. One limitation is that cell cluster labels are needed a *priori*. While we automatically use Celda to identify cell clusters if none are supplied, any fast cell clustering approach can be substituted. As the contamination distribution for each cell population is derived from all other populations present in the dataset, it may sometimes better to use broader cell population labels. For example, including all T-cells in one cluster rather than treating individual T-cell subpopulation as a separate subcluster may help alleviate T-cell specific counts in the calculation of the contamination distribution. Overall, computational decontamination of single-cell counts with DecontX will aide in down-stream clustering and visualization and can be systematically included in analysis workflows.

## ONLINE METHODS

### Statistical model

We assume there are *K* known distinct cell populations among the *M* cell samples, where cell *j* has *N*_*j*_ observed transcripts. We denote native expression distribution for cell population *k* as a G-length vector *ϕ*_*k*_. For the notational convenience, we will use *ϕ*_*−k*_ = {*ϕ*_*k*_’: *k*’ ≠ *k, k*’ ∈ {1, 2, …, *K*}} to represent gene expressions from all other cell populations other than *k*. Each cell *j* has a parameter *θ*_*j*_ to represent the proportion of counts that are derived from native expression distribution. *θ*_*j*_ is assumed to from a global beta distribution which leverages the variation of contamination level cross all the cells in the dataset, with hyperparameters *a*_1_ and *a*_2_ *a-priori*. The *t*^*th*^ transcript *x*_*jt*_ in cell *j* has a hidden state, *y*_*jt*_, which follows a bernoulli distribution parameterized by *θ*_*j*_ and denotes the transcript’s membership to be native expression distribution (*y*_*jt*_ = 1) or contamination distribution (*y*_*jt*_ = 0). Assuming that transcripts are conditionally independent given hidden state *y*_*jt*_ and cell’s population *z*_*j*_, *x*_*jt*_ follows a multinomial distribution either parameterized by 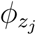 denoting native expression or 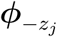 denoting contamination. The joint posterior distribution can be expressed as:

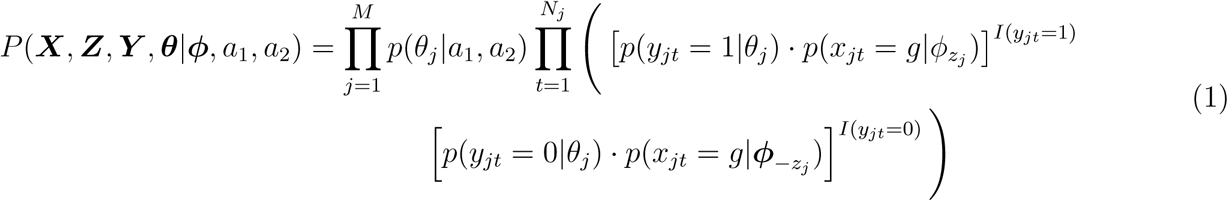

To simplify conputation work and notation, we assume the contamination distribution *η*_*k*_ is a simple linear combination of *ϕ*_*−k*_.

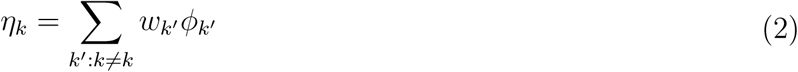

where the weight *w*_*k′*_ is the proportion of native transcripts from cluster *k*′ and is calculated using expected values, of which the full definition is given later in inference.

### Variational inference

We use variational inference to approximate the probability densities for our model. The following variational distributions are introduced to break down the coupling of ***θ*** and ***Y*** for variational inference:

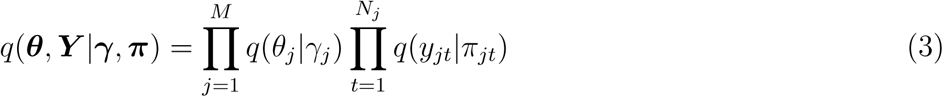

where the Beta parameter *γ*_*j*_ = {*γ*_*j*1_, *γ*_*j*2_}, and Bernoulli parameter *π*_*jt*_ = {*π*_*jt*1_,*π*_*jt*2_} are the free variational pameters. *π*_*jt*_ satisfies *π*_*jt*1_ + *π*_*jt*2_ = 1, and and 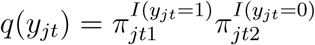. The variational Beta distribution for 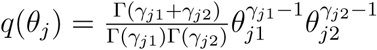

The need to compute the expectation of the *θ* _*j*_ arises in deriving the variational inference. Using the general fact for exponential family, that the derivative of the log normalization factor with respect to the natural parameter is equal to the expectation of the sufficient statistic (log *θ*_*ji*_, *i* ∈ {1, 2} in our Beta distribution), we have:

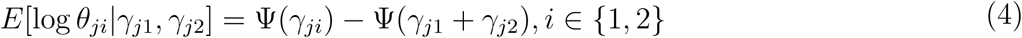

where Ψ is the digamma function, the first derivative of the log Gamma function. For simplicity in notation, let us use *Q* = {***θ***, ***Y***} and *a* = {*a*_1_, *a*_2_}. We begin variational inference by bounding the log-likelihood using Jensens inequality.

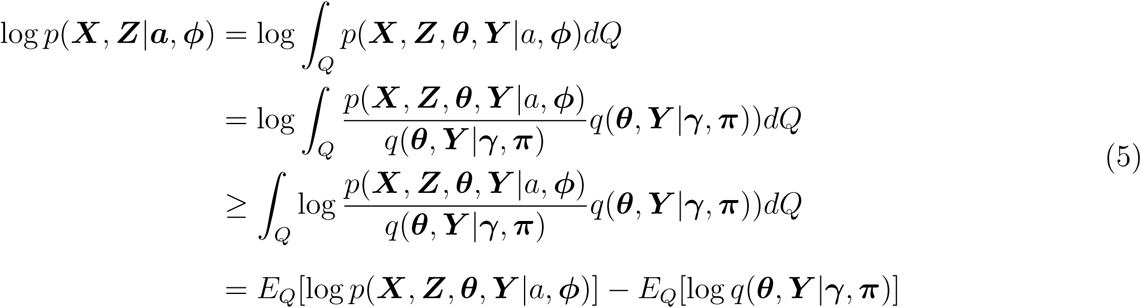

Jensens inequality provides us with a lower bound on the log likelihood for an arbitrary variational distribution *q*(***θ***, ***Y*** |*γ*, ***π***).

We then expand the lower bound:

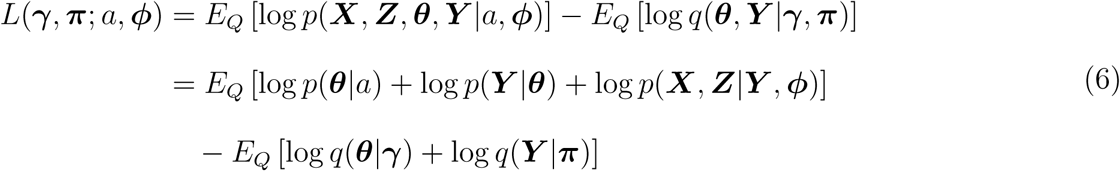

Expanding each term in the lower bound by taking expectation with respect to (***θ***, ***Y*** |*γ*, ***π***):

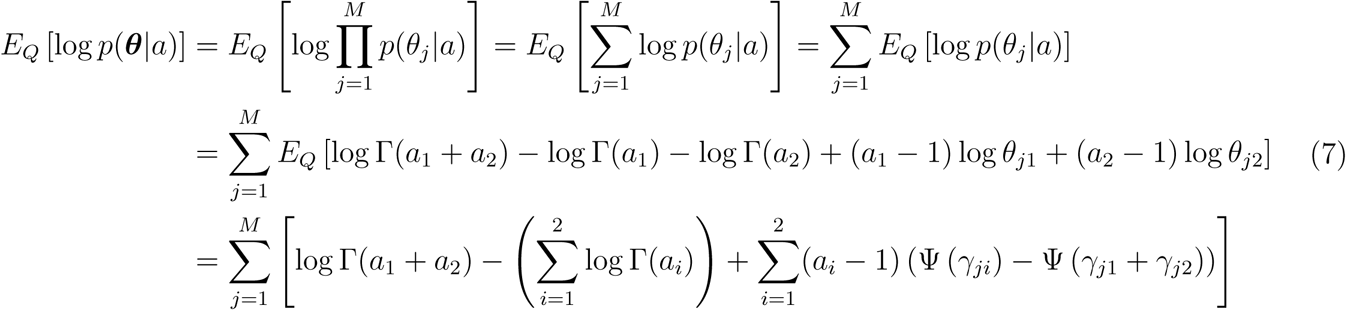

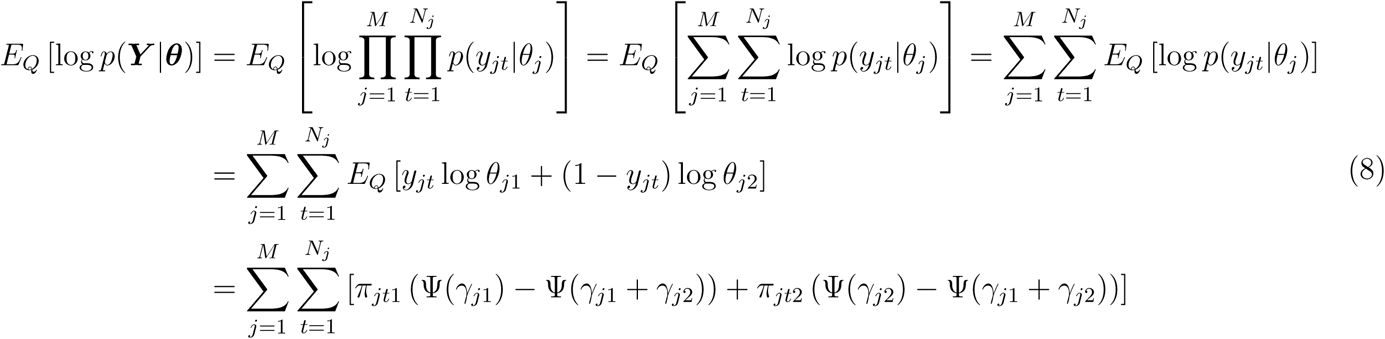

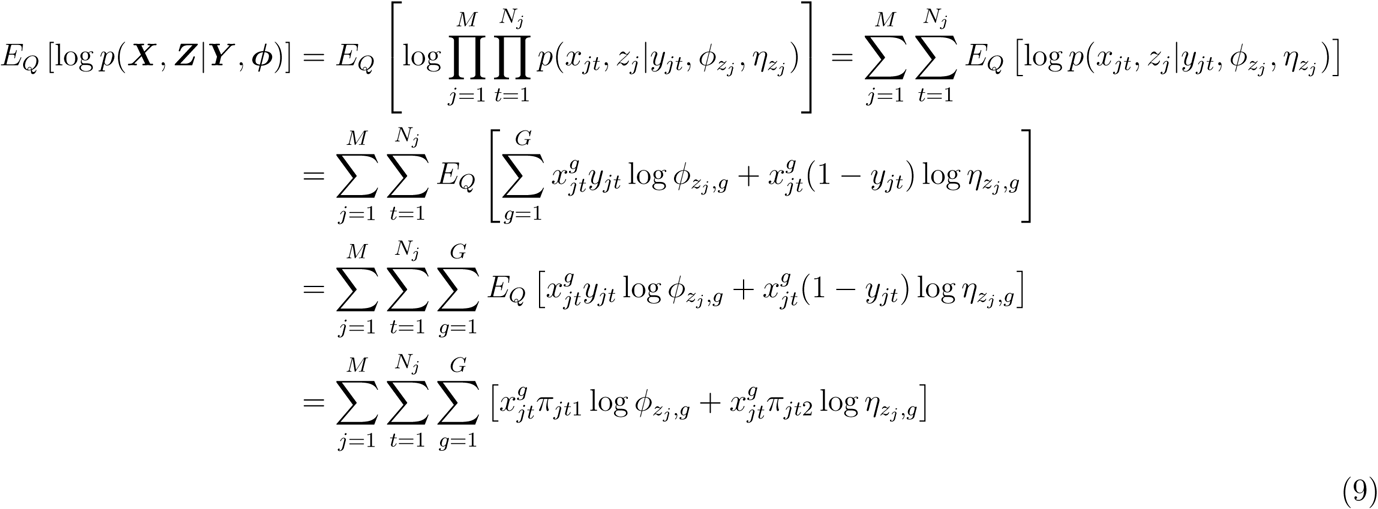

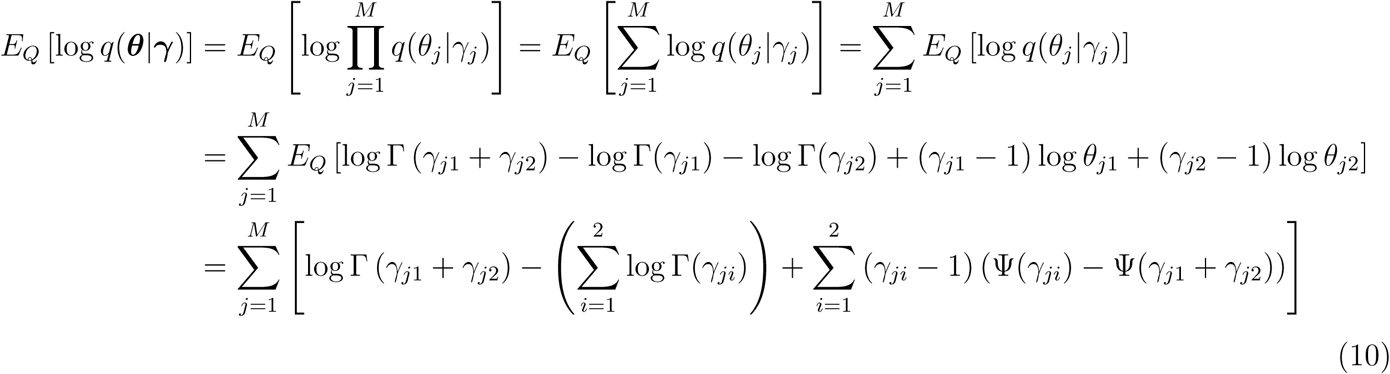

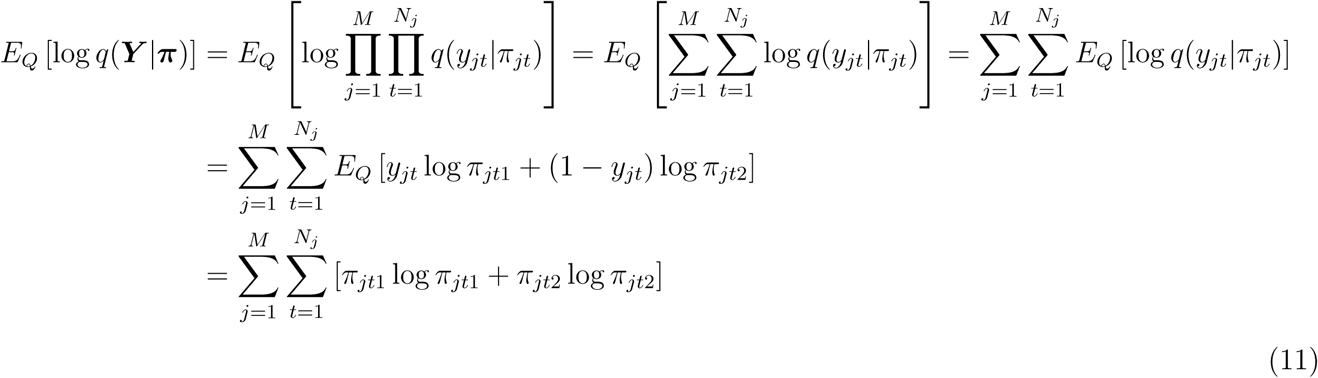

We then maximize the lower bound with respect to the variational parameters ***γ*** and ***π***.

First to maximize the lower bound with respect to ***π***. Since *π*_*jt*_ are independent, for *t* ∈ {1, 2, …, *N*_*j*_}, we isolate the terms that contains *π*_*jt*_. Lagrangian multiplier is added due to the constraint *π*_*jt*1_ + *π*_*jt*2_ = 1. We substituted 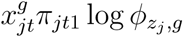 and 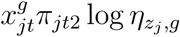 from equation (9) with 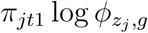 and 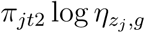 respectively, since 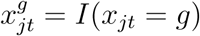 and is observed

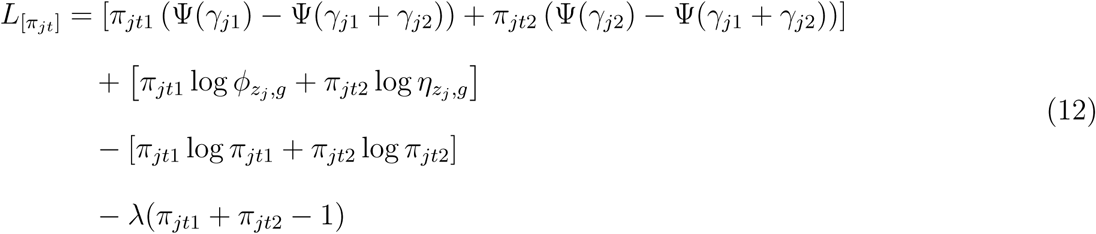

Taking derivative with respect to *π*_*jt*1_, we obtain:

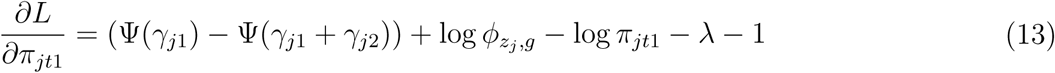

Setting this derivative to zero yields the maximizing value of the variational parameter *π*_*jt*1_:

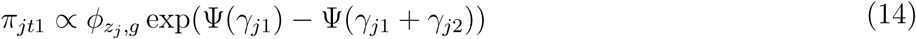

Similarly we could have *π*_*jt*2_:

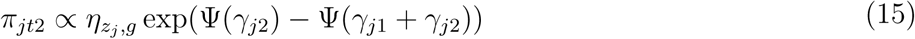

Next, we maximize the lower bound with respect to ***γ***. Since *γ*_*j*_ are independent for *j* ∈ 1, 2, …, *M*, each *1*_*j*_ can be estimated separately. We isolate the terms that contain *γ*_*j*_.

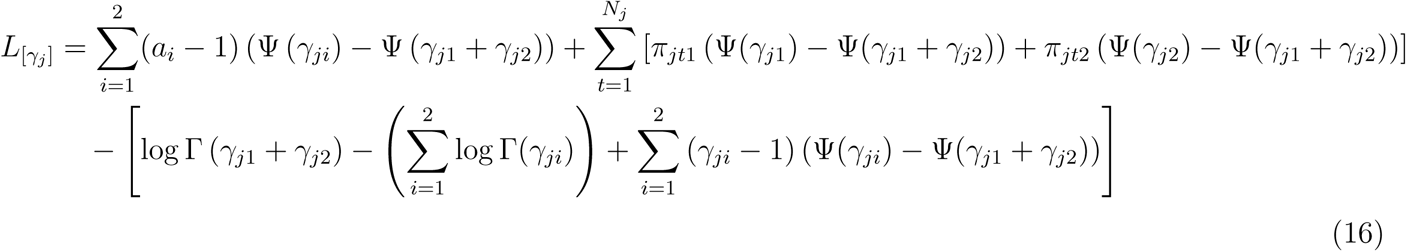

Taking derivative with respect to *γ*_*ji*_, we obtain:

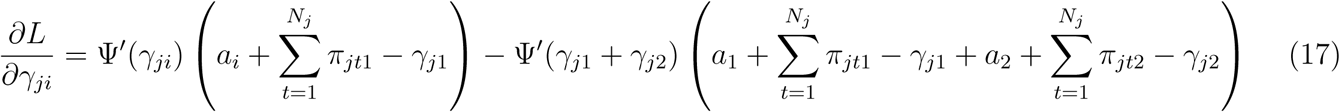

Where Ψ′ is the derivative of the digamma function. Setting this derivative to zero yields a maximum at:

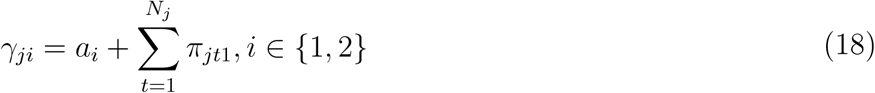

Finally, we move forward to estimating *ϕ, a*, and to update ***η***.

To maximize with respect to *ϕ*_*k*_, we isolate terms and add Lagrangian multiplier due to the constraint 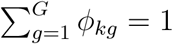:

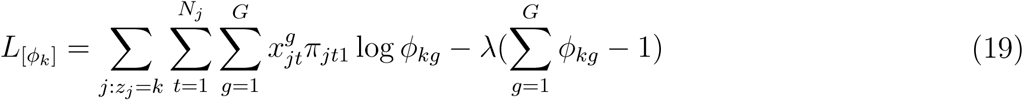

Taking the derivative with respect to *ϕ*_*kg*_ and set it to zero, we get:

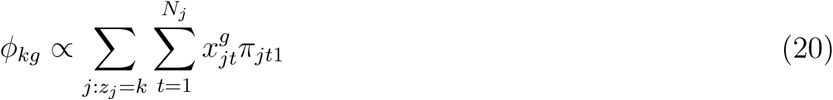

The weight *w*_*k*′_ is the proportion of native transcripts from cluster *k*′ and is calculated using expected values:

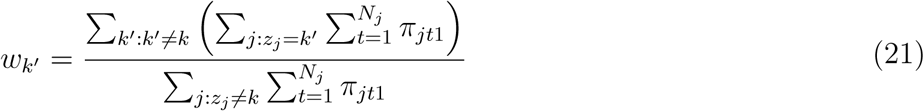

Hence we have our updated 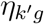 as:

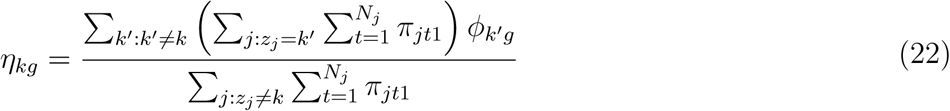

To maximize with respect to *a*, we isolate terms and get:

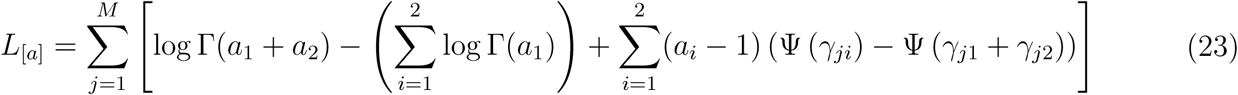

A Newton iteration can be used to find the maximal point *a* (Minka,2000), *which requires both the first and second derivatives of L*_[*a*]_. The first derivative, gradient *g*, and the second derivative, Hessian matrix *H* are:

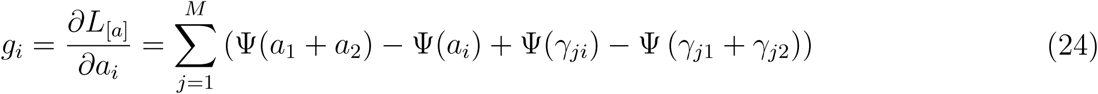

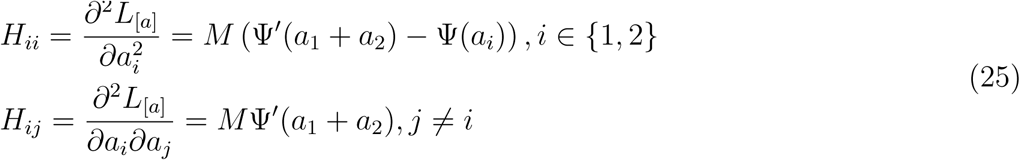

One Newton step is then

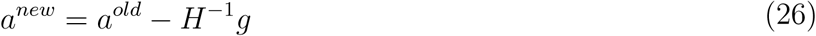

### Analysis of sorted human-mouse mixture single-cell dataset

A mixture of fresh frozen human (HEK293T) and mouse (NIH3T3) cells were sequenced together in 10X Genomics Chromium. This data is available at https://support.10xgenomics.com/single-cell-gene-expression/datasets/2.1.0/hgmm_12k. 6,164 human cells, 5,915 mouse cells and 741 multiplets were detected by CellRanger. Excluding multiplets, 12,079 cells with CellRanger predicted cell type were used to estimate contamination using DecontX.

### Analysis of sorted PBMC single-cell datasets

9 publicly-available PBMC datasets totalling of 84,432 cells were obtained from 10X Genomics. Each dataset consisted of a population of cells that were isolated with flow cytometry based on expression of a predefined protein marker. Cell populations included progenitor cells(CD34+), monocytes (CD14+), B cells (CD19+), Natural Killer cells (CD56+), helper T-cells (CD4+), regulatory T-cells (CD4+/CD25+), native T-cells (CD4+/CD45RA+/CD25-), naive cytotoxic T-cells (CD8+/CD45RA+) and cytotoxic T-cells (CD8+). A total of 7,363 genes which contained at least 3 counts across 3 cells were included in the analysis. DecontX used cell label by flow cytometry to estiamte contamination. Celda was used to identify and 76 gene modules and 21 cell clusters, including 8 clusters predominantly expressing T-cell markers, 2 clusters predominantly expressing Natural Killer cell markers, 2 clusters predominantly expressing B-cell markers, 2 clusters predominantly expressing monocyte markers, and 7 clusters predominantly expressing CD34 progenitor cell markers. These computationally inferred cell type labels were used in downstream analyses that examined the percentage of cells that express various marker genes. Using computationally derived cell clustered mitigated instances where a cell was improperly sorted and labeled by flow cytometry as belonging to one population when in fact it transcriptionally similar to another population.

### Analysis of the 4K PBMC single-cell dataset

4,340 PBMCs from a healthy donor were sequenced in a single channel of the 10X Genomics Chromium. Data is available at https://support.10xgenomics.com/single-cell-gene-expression/datasets/2.1.0/pbmc4k. A total of 4,529 genes which contained at least 3 counts across 3 cells were included in the analysis. 19 cell clusters and 150 gene modules were identified with Celda. Cell clusters 2 and 3 were classified as B-cells (MS4A1+); cell clusters 5, 6, 7, 8, 9 and 11 were classified as T-cells (CD3D+/CD3E+); cell clusters 13 and 14 were identified as LYZ+ monocyte group; cell cluster 15 was identified as FCGR3A+ monocytes group; cell cluster 10 was identified as NKG7+ and GNLY+ NK-cell group; cell clusters 18 and 19 were identified as FCER1A+ dendritic cell group; cell cluster 4 was identified as IRF7+ and IRF8+ plasmacytoid dendritic cell group; cell cluster 16 was identified as PPBP+ Megakaryocytes; cell cluster 1 was identified as IGHG1+ and IGHG2+ plasma cell group; cell cluster 12 was identified as CD34+ cell group; cell cluster 17 is likely to be multiplets for it has shown IL7R, CD3D and CD14 markers. DecontX used Celda estimated cluster label to estimate contamination.

## Supporting information

Supplementary figures and their legends

## ACKNOWLEDGMENTS

This work was funded by LUNGevity Career Development Award (J.D.C.) and Informatics Technology for Cancer Research (ITCR) 1U01 CA220413-01 (W.E.J.). We thank Carter Merenstein, Ke Xu and Xinyi Shi for helpful suggestions during the analysis.

## AUTHOR CONTRIBUTIONS

JDC coneived the project; JDC, SY, MY developped the model; SY, YK performed the analysis; SY, JDC, MY wrote the manuscript; SC, ZW assisted in software development; SY, JDC, MY, YK, SC, ZW, ZW, EJ reviewed the manuscript.

