## Supplementary figures and their legends for "Decontamination of ambient RNA in single-cell RNA-seq with DecontX"

### Supplementary Figure 1

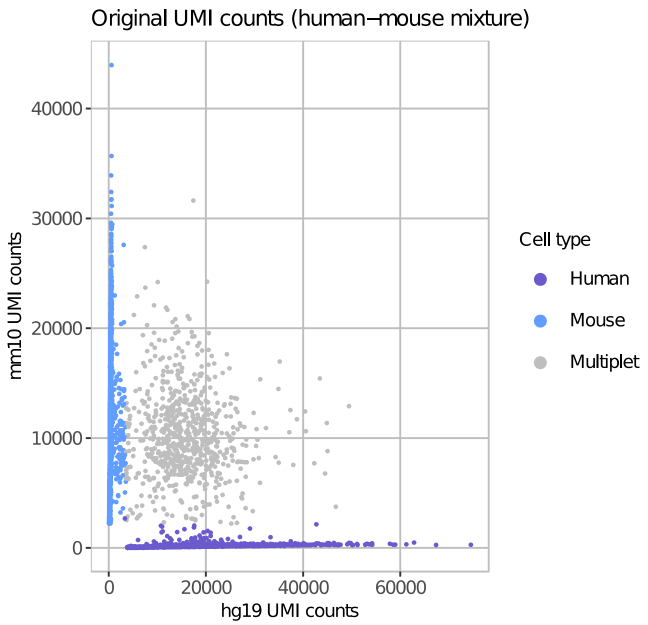


**Supplementary Figure 1: Identification human and mouse cells in droplets.** The number of UMIs aligned specifically to mouse genome is plotted against to the number of UMIs aligned specifically to human genome for each droplet. CellRanger was used to predict whether each droplet contained a human cell (purple), mouse cell (blue), or multiplet (grey). The multiplets were excluded from down-stream decontamination analysis.

### Supplementary Figure 2

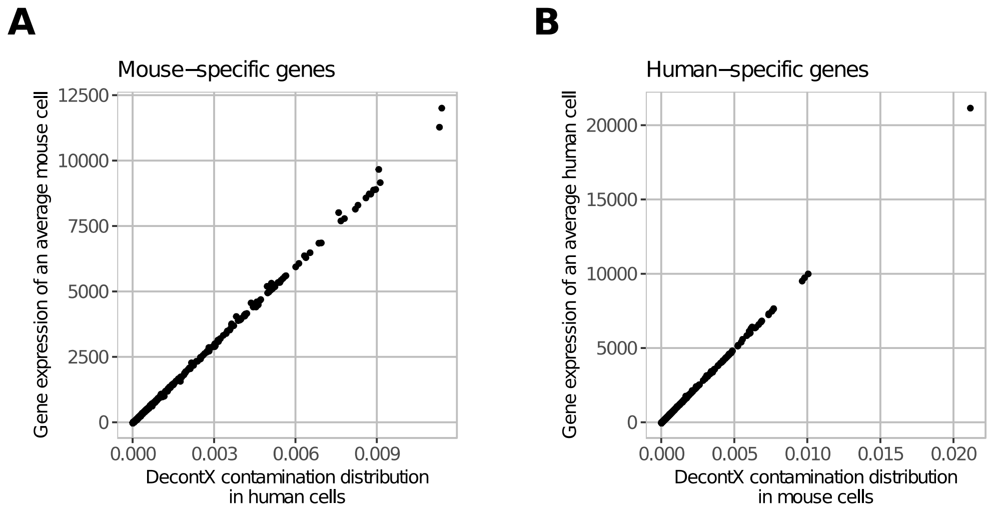


**Supplementary Figure 2: Comparisons between distributions of population-specific contamination and native expression in the mouse-human mixture dataset. (A)** A high correlation was observed between the gene probabilities in the DecontX-estimated contamination distribution for the human cell population and the gene expression levels within an average mouse cell. Each point represents a gene in the mouse transcriptome. **(B)** A high correlation was observed between the gene probabilities in the DecontX-estimated contamination distribution for the mouse cell population and the gene expression levels within an average human cell. Each point represents a gene in the human transcriptome.

### Supplementary Figure 3

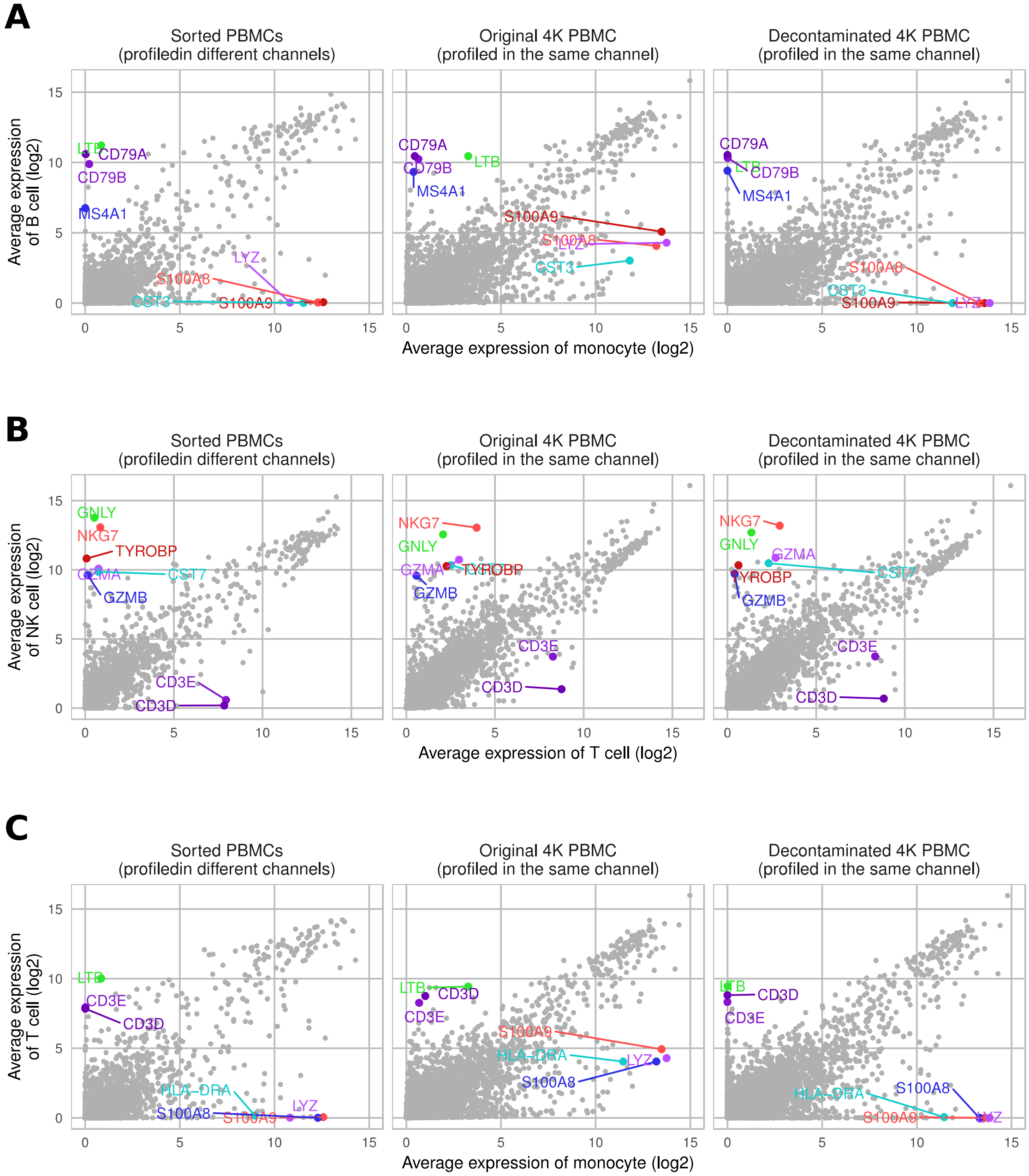

### Supplementary Figure 3

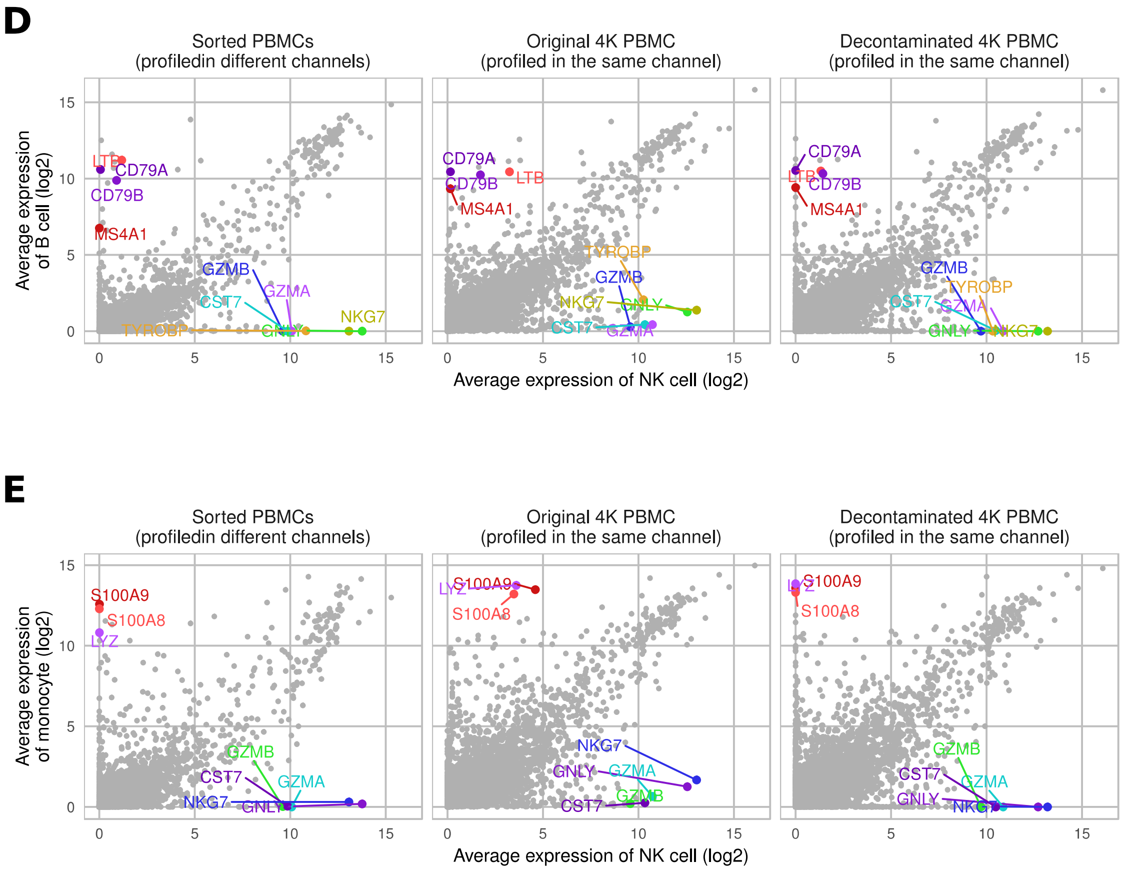


**Supplementary Figure 3: Expression of PBMC cell-type specific marker genes across cell populations.** For each gene, the average expression across all cells in a population is plotted against the average expression across all cells in another population. **Left:** data from sorted PMBCs profiled in different channels. **Middle:** data from the PBMC 4K before decontamination with DecontX; **Right:** PBMC 4K data after decontamination with DecontX. Comparisons of cell populations include: **(A)** B-cells vs monocytes, **(B)** NK-cells vs T-cells, **(C)** T-cells vs monocytes, **(D)** B-cells vs NK-cells, and **(E)** monocytes vs NK-cells.

### Supplementary Figure 4

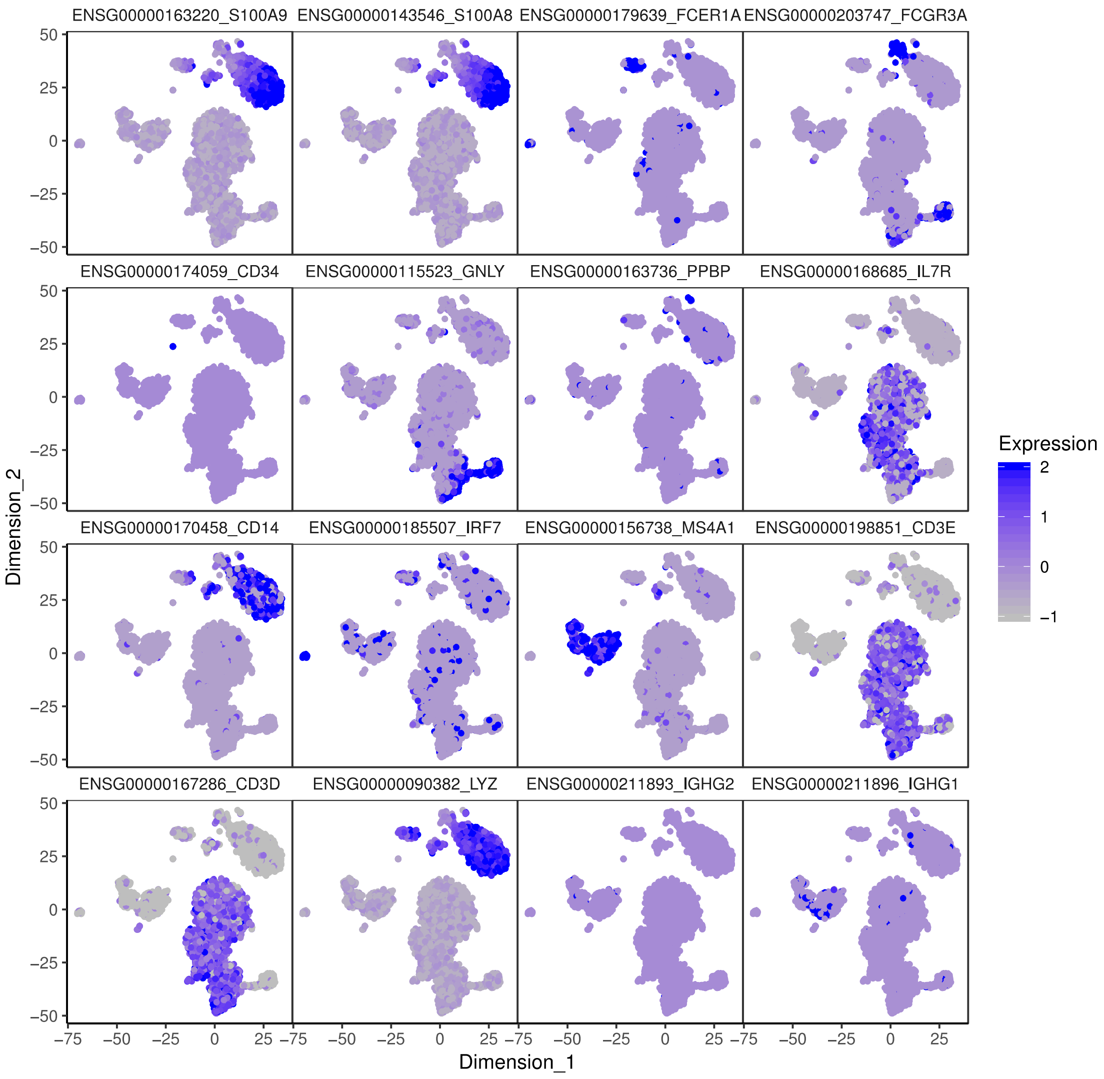


**Supplementary Figure 4: Expression of PBMC markers in the 4K dataset.** A tSNE was generated using the module probabilities derived from Celda. Each point is a cell and is colored by the relative expression level of the specific marker gene using the original 4k PBMC dataset before decontamination.

### Supplementary Figure 5

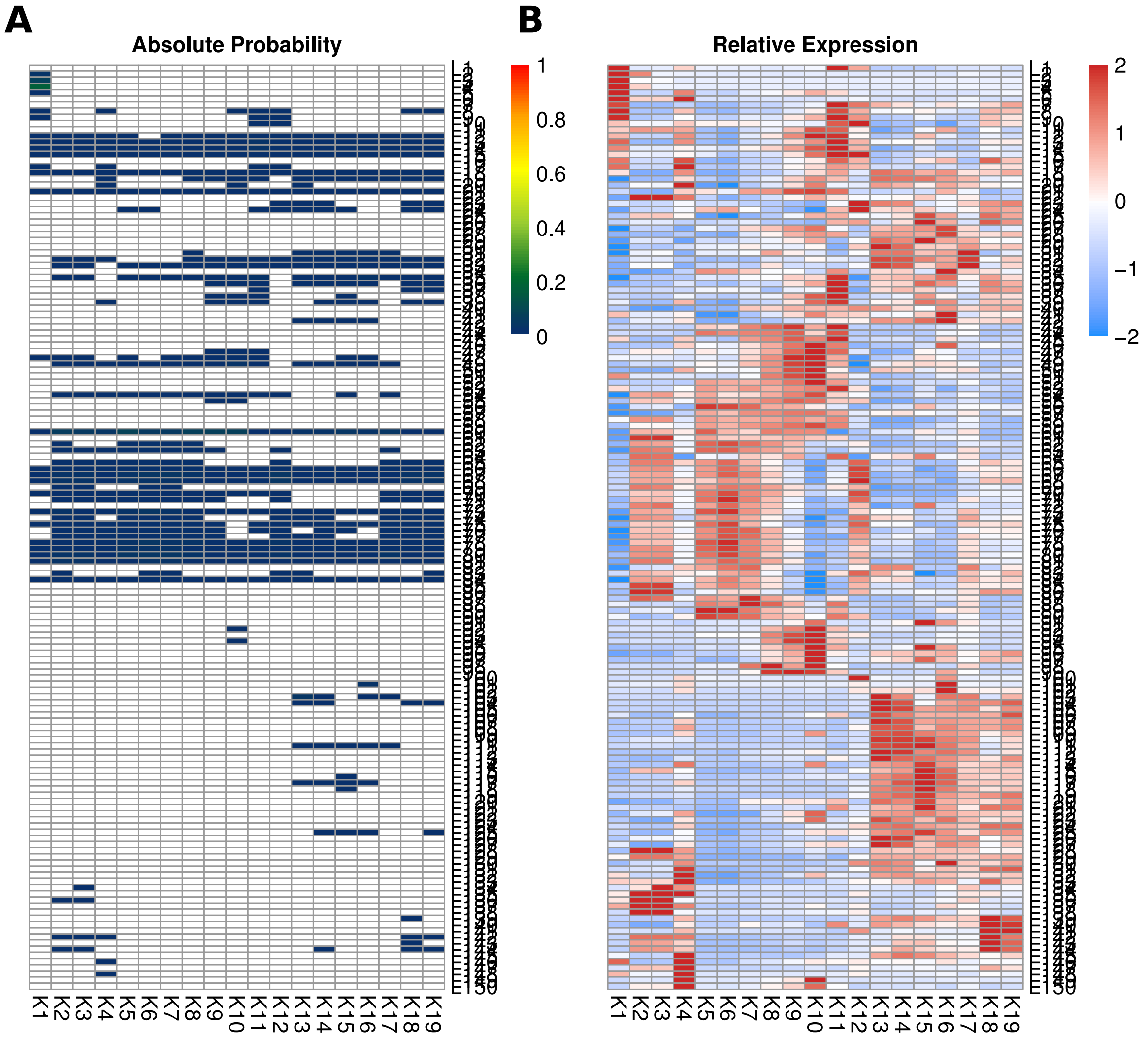


**Supplementary Figure 5: Celda probability and relative expression heatmaps for PBMC 4K. (A)** The probability for each of 150 modules (rows) is shown in each of the 19 cell populations (columns). **(B)** The relative expression for each of 150 module (rows) is shown for each of the 19 cell populations (columns).
